# Discovery of molecular glues by modeling ternary complex conformational ensembles and thermodynamic stability

**DOI:** 10.1101/2025.01.13.632817

**Authors:** Jesus A Izaguirre, Yujie Wu, Huafeng Xu, Zachary McDargh, Fabio Trovato, Timothy Palpant, Asghar M Razavi, Cheryl Koh

## Abstract

The rational design of molecular glue degraders is challenging because glue-mediated protein degradation depends on a complex interplay of molecular mechanisms: an effective molecular glue must simultaneously bring together two distinct proteins to form an interface that is both stable and accessible for ubiquitination by E3 ligases. We present GlueMap, a computational platform for discovering and optimizing molecular glues (MGs) through integrative structural and pharmacodynamic modeling. GlueMap combines molecular dynamics (MD) simulations and machine learning (ML) to model targeted protein degradation (TPD) mediated by molecular glues. We validated GlueMap by applying it to two case studies: DDB1-dependent degradation of CDK12 and CRBN-dependent degradation of GSPT1. In the DDB1–CDK12 system, GlueMap revealed strong correlations between ternary-complex stability, quantified through rigorous free-energy calculations, and experimental degradation efficiency. Notable outliers underscored the importance of incorporating conformational dynamics within assembled Cullin–RING ligase complexes. Using a supervised variational autoencoder combined with attention-based regression, GlueMap achieved high predictive accuracy for degradation potency from structural features. For CRBN–GSPT1, GlueMap reproduced known ternary-complex crystal structures and identified previously unrecognized protein-protein interactions (PPIs) that may guide the design of next-generation glues. Hierarchical virtual screening of more than 12 billion compounds successfully recovered known degraders and uncovered nine new potent MGs with DC_50_ values below 50 nM. Across both systems, GlueMap outperformed Boltz-1 and Boltz-2, state-of-the-art structure-prediction models that failed to reproduce the CDK12–DDB1 crystal structure and captured substantially less PPI diversity. By integrating structural, thermodynamic, and ubiquitination metrics, GlueMap establishes a multi-criteria framework for rational molecular glue design. These results demonstrate that GlueMap provides both mechanistic insight and practical acceleration of molecular glue discovery and optimization across diverse therapeutic targets.

## Introduction

Molecular glues (MGs) are therapeutic small molecules that overcome key limitations of heterobifunctional degraders, owing to their smaller size and potentially superior physicochemical properties.^1^ Most MG discovery efforts rely on phenotypic screening, followed by target identification and mechanism-of-action validation.^2,3^

The computationally enabled rational design of MG degraders follows two primary approaches: (1) predicting how a candidate MG modifies protein surfaces to induce specific protein–protein interactions (PPIs), and (2) predicting PPIs that form in solution and evaluating how MGs stabilize the resulting ternary complexes. We follow the latter approach. Several resources provide comprehensive background on molecular glue biology and PPI prediction. Huan *et al.*^4^ offer an in-depth review of weak PPI identification (encounter complexes); Kozicka *et al.*^5^ present a curated dataset of MG degraders; and Schreiber^6^ discusses induced proximity degraders. Konstantinidou and Arkin^7^ provide a recent review of MGs discovered from 2014 and 2023. The MGTBind database catalogs molecular glue bioactivity data and ternary interactions.^8^

Recent advancements in machine learning (ML) have transformed the field. AlphaFold2^9^ revolutionized protein structure prediction and enabled new approaches to modeling proteinprotein complexes.^10^ Building on this foundation, AlphaFold3^11^ extended these capabilities to biomolecular interaction prediction, including protein–ligand and ternary complexes. Multiple open-source alternatives have rapidly emerged, including Boltz-1,^12^ Boltz-2,^13^ Chai-1,^14^ and RoseTTAFold All-Atom.^15^ Prael *et al.*^16^ developed methods to predict CRBN-based molecular glues by screening large compound libraries. Monte Rosa scientists^17^ computationally mined the human proteome to predict more than 1,400 CRBN-degradable targets. More recent developments suggest potential integration with advanced models like AlphaFold3 or Boltz-1 to enhance predictive accuracy. A preprint^18^ introduced an approach similar to these models for predicting MG-mediated ternary complexes.

Despite these advances, recent benchmarks reveal significant limitations in ML-based methods for predicting molecular glue-mediated ternary complexes. Comprehensive evaluations^19^ using curated datasets with 88–221 ternary complex structures show that AlphaFold3 achieves only 33–51% success rates for recovering molecular glue-protein interactions, with many correct predictions likely derived from training data memorization rather than generalization. Systematic studies^20,21^ demonstrate that ML models struggle particularly with shallow PPIs characteristic of MG-induced complexes, where AlphaFold-Multimer shows intrinsic limitations. The CASP15 assessment^22,23^ demonstrated that while protein assembly prediction success rates reached 90% for evolutionarily related complexes, accuracy was much lower (20–30 percent) for antibody–antigen and nanobody interactions, a scenario analogous to many induced proximity systems. Additional benchmarks^24,25^ reveal that even when AlphaFold3 produces structurally accurate protein-protein complexes, single-seed predictions exhibit 60% failure rates, and binding free-energy estimates from ML-predicted structures are less accurate than those derived from experimental templates.

These persistent challenges, particularly for novel protein-protein interfaces, conformational flexibility, and induced proximity complexes, motivate complementary computational approaches that integrate physics-based simulations and thermodynamic modeling. Our approach utilizes physics-based simulations to model conformational landscapes and design MGs that can stabilize weak basal interactions.^26^ The framework integrates multiple components, including machine learning models that demonstrate high accuracy in predicting degradation potency from structural features. Key elements include encounter and ternary complex prediction, molecular dynamics (MD) simulations for generating conformational landscapes, analyzed through Markov^27^ and Non-Markov State models,^28^ binding affinity predictions for binary and ternary complexes, ubiquitin-lysine distance predictions, and neural network models to derive experimental observables from MD data. These tools are integrated into GlueMap, a molecular glue discovery framework that builds on our prior work.^29,30^

The computational framework for molecular glue discovery has matured significantly, with emerging specialized tools^31–33^ and integrated platforms^34,35^ demonstrating the value of combining structure prediction, molecular dynamics, free energy calculations, and machine learning. Alternative computational approaches that primarily rely on docking^36–38^ or simplified modeling can serve as starting points for our more comprehensive approach.^39^ Reviews of current discovery strategies^40–42^ highlight the complementary roles of computational and experimental methods, with physics-based approaches offering advantages for systems where ML methods show limitations.^43^

We consider two case studies: the DDB1-mediated molecular glue degradation of CDK12 and Cyclin K, for which there is abundant data for validation, and the CRBN-mediated molecular glue degradation of GSPT1, for which we designed novel molecular glues. GSPT1, a translation termination factor, plays a critical role in various cancers, including acute myeloid leukemia (AML) and MYC-driven solid tumors. Targeting GSPT1 for degradation has emerged as a promising therapeutic strategy, particularly through MGs that recruit CRBN to facilitate GSPT1 degradation.^44^ GSPT1 degradation induces TP53-independent cell death while sparing normal hematopoietic stem cells,^45^ making it a viable strategy for selective cancer therapy.

### Overview of GlueMap Workflow

In GlueMap (Fig. 1), we integrate protein structure prediction, protein-protein docking, and advanced simulations to predict structural ensembles of encounter complexes between POIs and effector proteins, as well as ternary complexes involving POIs, molecular glues, and effector proteins.

**Figure 1:**
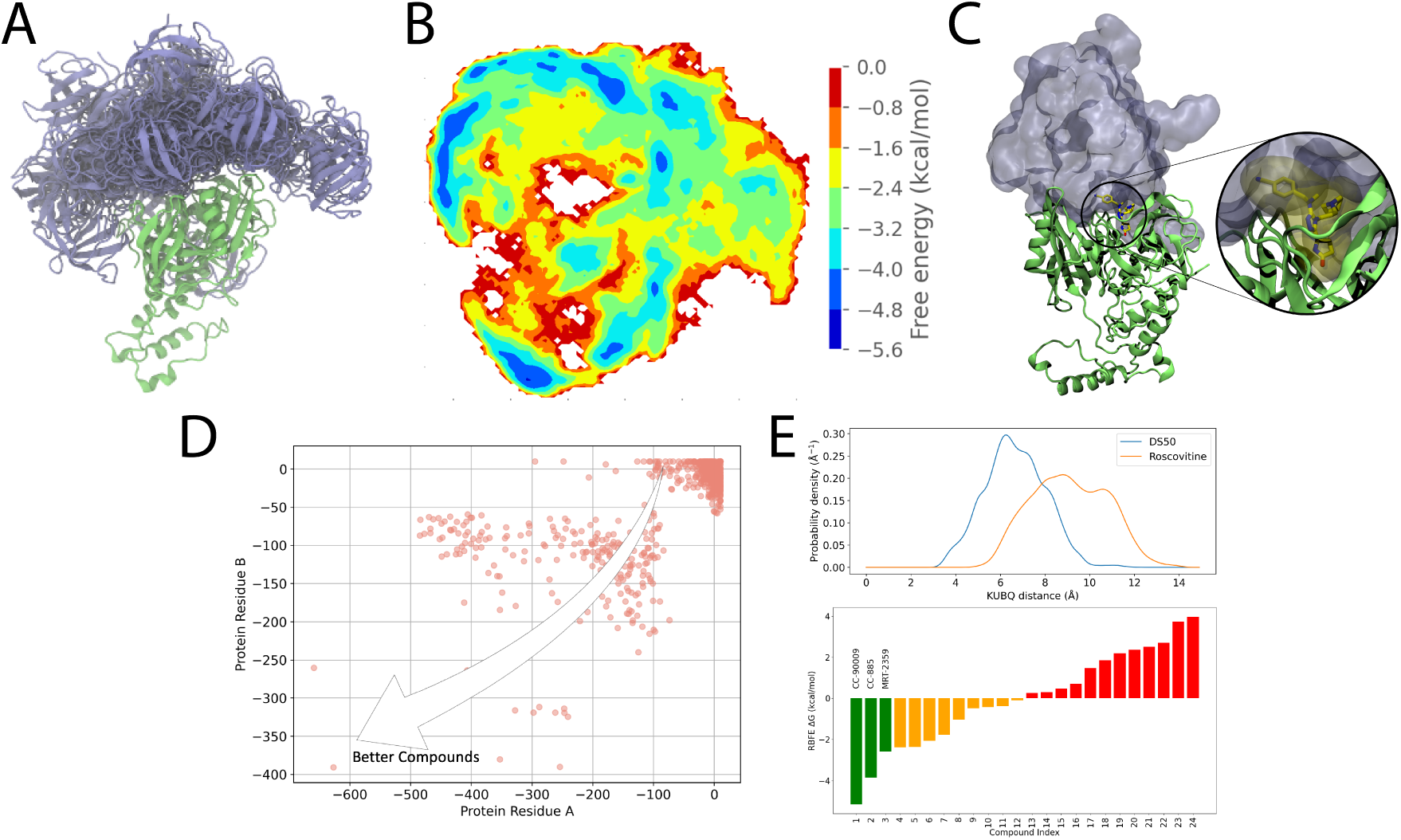
Overview of the computational process for **GlueMap**: (A) Protein-Protein interface (PPI) docking followed by long MD simulations. (B) Construction of a PPI free energy landscape. (C) Small molecule virtual screening against PPIs of interest from the conformational ensemble. (D) Initial Compound selection for longer MD simulations using scoring function. (E) Final compound selection, using structural stability of the ligand in the pocket, ubiquitination profiles and ternary complex stabilities (binding free energies).

The workflow begins with protein-protein docking to identify plausible interaction surfaces. While we use MOE’s protein-protein docking tool,^46^ alternative methods can also be employed. Following docking, all-atom MD simulations (1 to 2 µs) are conducted for each predicted PPI. These simulations refine the interaction interfaces using physics-based force fields, providing a more accurate representation of the molecular interactions (Fig. 1a).

Next, a PPI free energy landscape is constructed by integrating the MD simulation data using Markov State Modeling (MSM) and related techniques. This approach enables the identification and selection of metastable PPIs for further analysis (Fig. 1b). PPIs are selected based on interfacial pocket druggability and potential for ubiquitination.

We then perform small-molecule virtual screening against an ensemble of PPIs derived from the MSM. Pharmacophores are defined for each metastable PPI conformation to identify potential molecular glues, followed by rapid pharmacophore screening of virtual compound libraries, such as those available from Enamine (Fig. 1c).

To score the screened compounds, short MD simulations (10 ns each) are conducted on candidates filtered using pharmacophores. Electrostatic and van der Waals interactions are calculated for each protein residue, enabling the identification of key binding residues and pockets. Based on these results, compounds are selected for longer MD simulations (200 ns each) (Fig. 1d).

The final compound selection uses Pareto optimization, taking into account structural stability, thermodynamic stability—predicted by relative binding free energy (RBFE) calculations— and ubiquitination potential of the corresponding ternary complex to identify promising molecular glue candidates (Fig. 1e).

As demonstrated by applying GlueMap to various projects, the workflow should be tailored to meet the specific needs of each project. In some cases, only certain modules may be utilized, while in others, the generic steps outlined here may be replaced with more suitable alternatives. The modular design of GlueMap facilitates such customization.

## Results

### Case Study: DDB1-dependent CDK12-Cyclin K degradation

Kozicka *et al.*^5^ report a rich dataset of molecular glues that degrade CDK12 and Cyclin K through DDB1. This dataset contains crystal structures, experimental binary and ternary affinities, and degradation results. We use this dataset to assess various capabilities of GlueMap.

#### Prediction of ternary complex structures

We modeled ternary complexes between DDB1 and CDK12, mediated by analogs of the CDK12 inhibitor CR8 (see Table 1 for a summary of simulations used in this case study). We start from docked CDK12-DDB1 poses, run Hamiltonian Replica Exchange Molecular Dynamics (HREMD) starting from the resulting poses (see Methods), cluster the conformations sampled in the simulations, and choose cluster centers of the highest populated clusters as our models. The resulting models are in good agreement with the crystal structures, which are *not* used as input to the simulations (Fig. 2).

**Figure 2:**
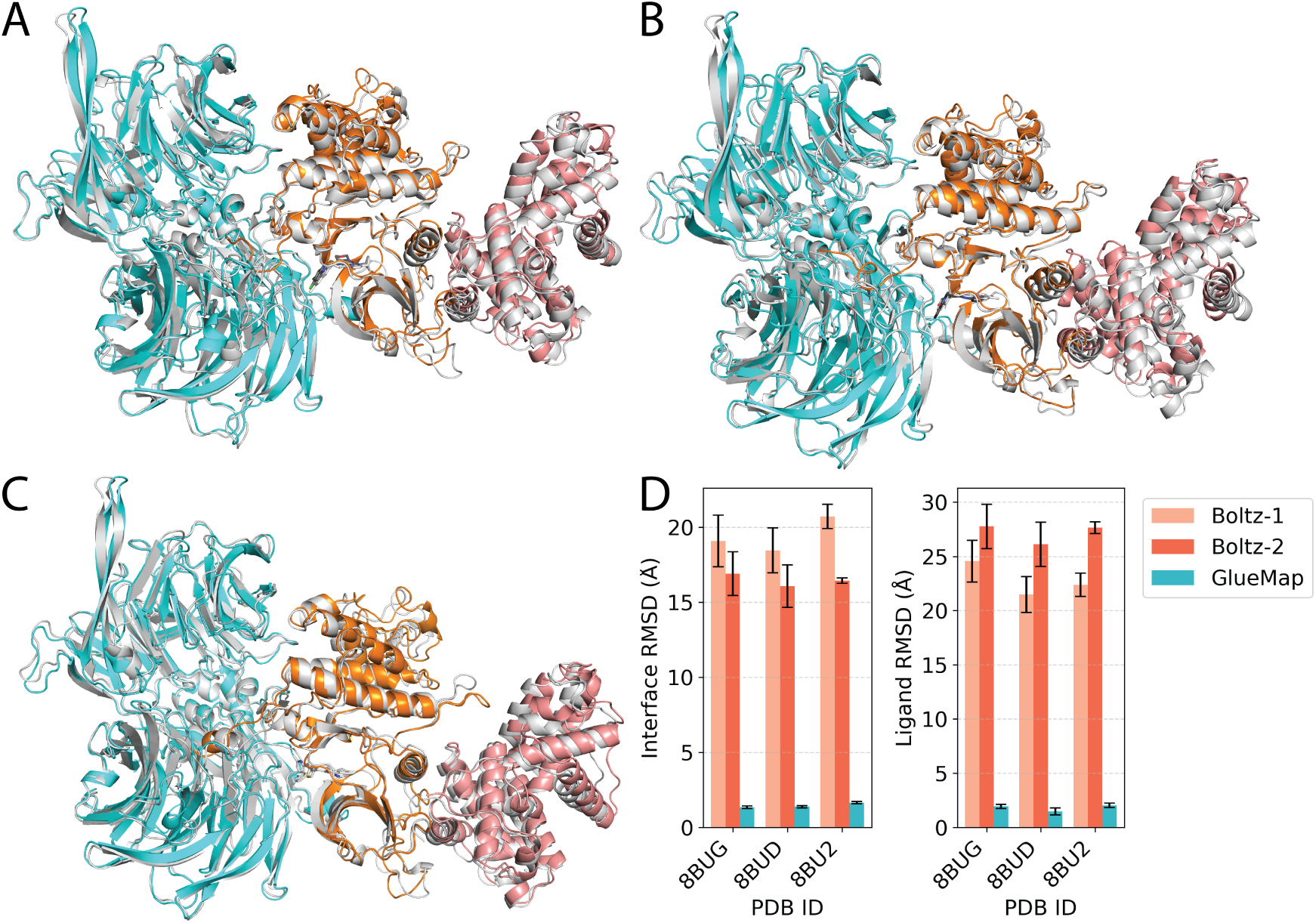
Comparison of modeled ternary complex structures with crystal structures for DDB1-CDK12-CYCK (A: PDB ID 8BU2; B: PDB ID 8BUD; C: PDB ID 8BUG). For each MG, the crystal structure belongs to the highest-population conformational cluster identified through GlueMap. (D) Comparison of the predictions of GlueMap to those of Boltz-1 and Boltz-2.

**Table 1:** Summary of the molecular simulations performed for the DDB1-CDK12-CycK case study. Tc = Ternary complex.

| Simulation type | Starting state | Time (μs) | Purpose |
| --- | --- | --- | --- |
| Protein-protein docking | – | – | Tc structure prediction |
| HREMD simulations | Docked poses | 2.9 | Tc structure prediction |
| HREMD simulations | Crystal structures | 16 | Tc conformational landscape |
| MD simulations | Crystal structures (superposed) | 0.1 | Ubiquitination prediction |

We compare our predictions to those of recent AI co-folding models Boltz-1 and Boltz2.^12,13^ We used each model to generate 20 structures of DDB1-MG-CDK12 ternary complexes, omitting CycK (see Methods). Both models failed to generate useful predictions of the ternary complex structures (Tables 2 and 3). This finding is consistent with recent systematic benchmarks of AI structure prediction models for predictions of ternary complexes and other complex systems.^19,47^

**Table 2:** Interface RMSD values of DDB1-MG-CDK12 ternary complex structure predictions from Boltz-1, Boltz-2, and GlueMap. The protein-protein interface was defined by selecting residue pairs in the crystal structure with at least one pair of heavy atoms within 10 Å. Using all heavy atoms from these residues, the predicted structures were superposed onto the crystal, and the heavy-atom RMSD was computed on the same atom set.

| PDB | Boltz-1 (Å) | Boltz-2 (Å) | GlueMap (Å) |
| --- | --- | --- | --- |
| 8BU2 | 20.7 ± 0.8 | 16.4 ± 0.2 | 1.6 ± 0.1 |
| 8BUD | 18.5 ± 1.5 | 16.1 ± 1.4 | 1.5 ± 0.1 |
| 8BUG | 19.1 ± 1.7 | 16.9 ± 1.5 | 1.3 ± 0.1 |

**Table 3:** Ligand RMSD values of structure predictions from Boltz-1, Boltz-2, and GlueMap. The ligand pocket was defined by selecting residues in the crystal structure with at least one heavy atoms within 10 Å of the ligand. Using all heavy atoms from these residues, the predicted structures were superposed onto the crystal, and the RMSD was computed on the ligand heavy atoms.

| PDB | Boltz-1 (Å) | Boltz-2 (Å) | GlueMap (Å) |
| --- | --- | --- | --- |
| 8BU2 | 22.4 ± 1.1 | 27.6 ± 0.5 | 2.3 ± 0.3 |
| 8BUD | 21.5 ± 1.7 | 26.1 ± 2.0 | 1.8 ± 0.4 |
| 8BUG | 24.6 ± 1.9 | 27.8 ± 2.0 | 2.3 ± 0.2 |

#### Predicting stability of DDB1-MG-CDK12 ternary complexes using relative binding free energy calculations

Ternary complex formation mediated by molecular glues consists of the following reactions

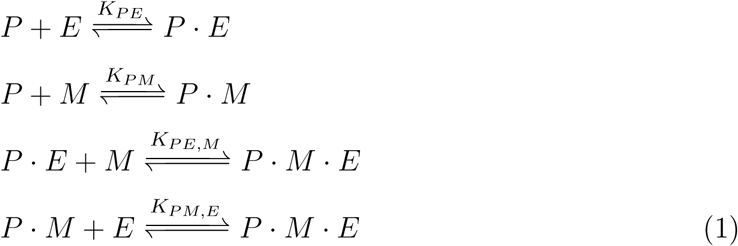

In this model, the molecular glue (M) is assumed to bind more strongly to the target protein P than to the E3 ligase E.^5,48^ For the DDB1-CDK12 system, the molecular glue specifically binds to CDK12 (P) but not to DDB1 (E).

Thermodynamic cycle closure implies

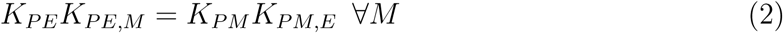

The ternary complex stability is governed by the equilibrium constants in Eq. 1. For a series of molecular glues, we can predict the binding constants *K_PM_* and *K_PE,M_* using relative binding free energy (RBFE) calculations.^49–52^ Together with the protein-protein binding constant *K_PE_*, determined from fitting to experimental titration data, we can quantitatively predict the ternary complex stability at given molecular glue concentrations.

Here we use RBFE to predict the thermodynamic stabilities of the binary complexes of MG with CDK12 and of the ternary complexes (Fig. 3) of MG, DDB1 and CDK12, obtaining results in good agreement with experimental values^5^ (see also Methods and Supplementary Information).

**Figure 3:**
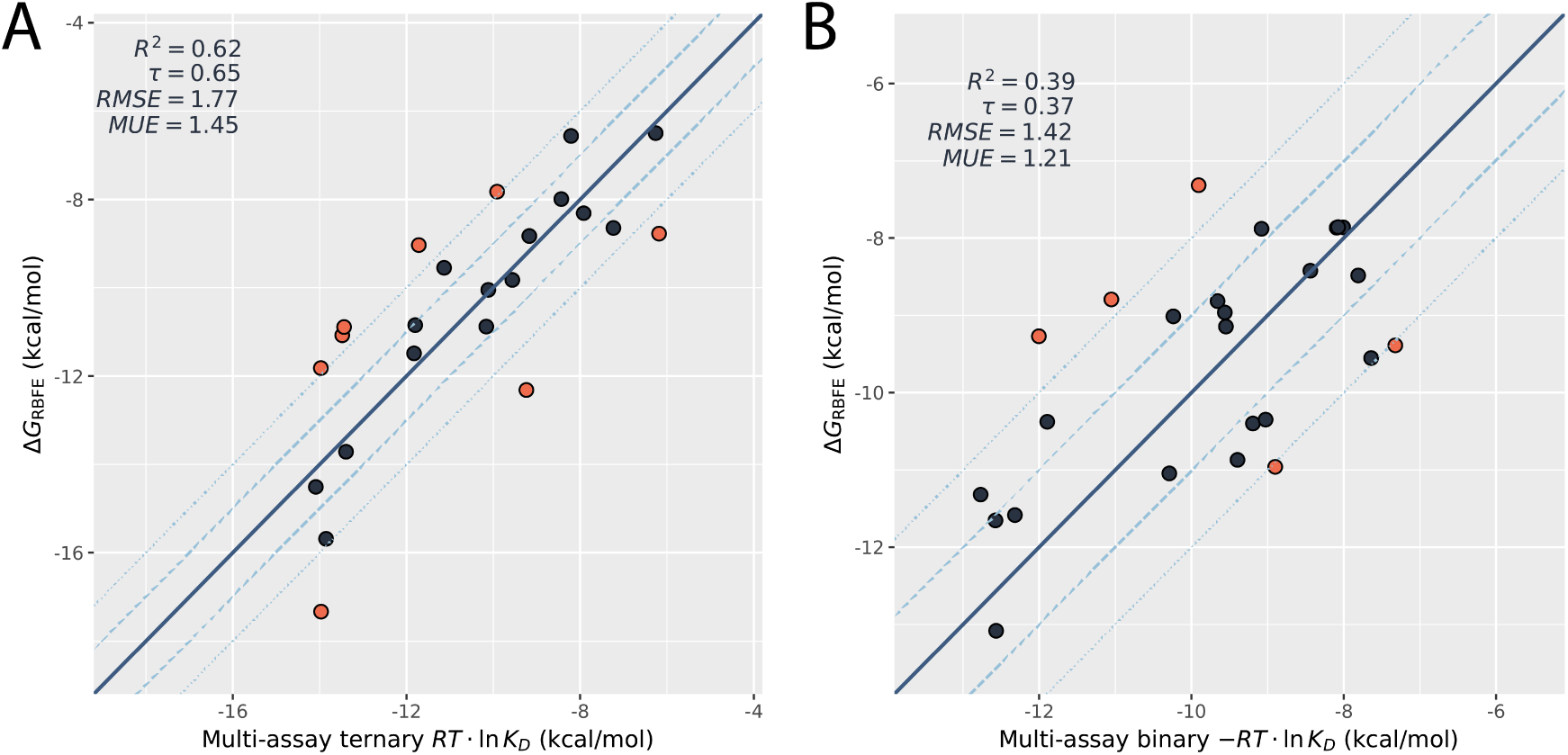
The predicted binding free energies for binary and ternary complexes compared to the experimental values for DDB1-CDK12. The experimental values of 10 glues (DS09, DS13, DS14, DS57, DS70, Z10, Z11, dCeMM2, dCeMM3, 919278, flavopiridol) are used as references to calibrate the DiffNet analysis; these reference glues are not included here. (A) The predicted binding free energies for binary complexes of molecular glues binding to CDK12. (B) The predicted binding free energies for ternary complexes of molecular glues binding to DDB1-CDK12.

The ternary complex stability can be estimated from these binary and ternary *K_D_*’s. In binary binding, which can be characterized with a single metric IC_50_, as the ligand concentration increases, the concentration of the binary complex approaches that of the receptor. In ternary complex formation, however, the ternary complex concentration reaches a different maximum for different molecular glues (Fig. 4a).

**Figure 4:**
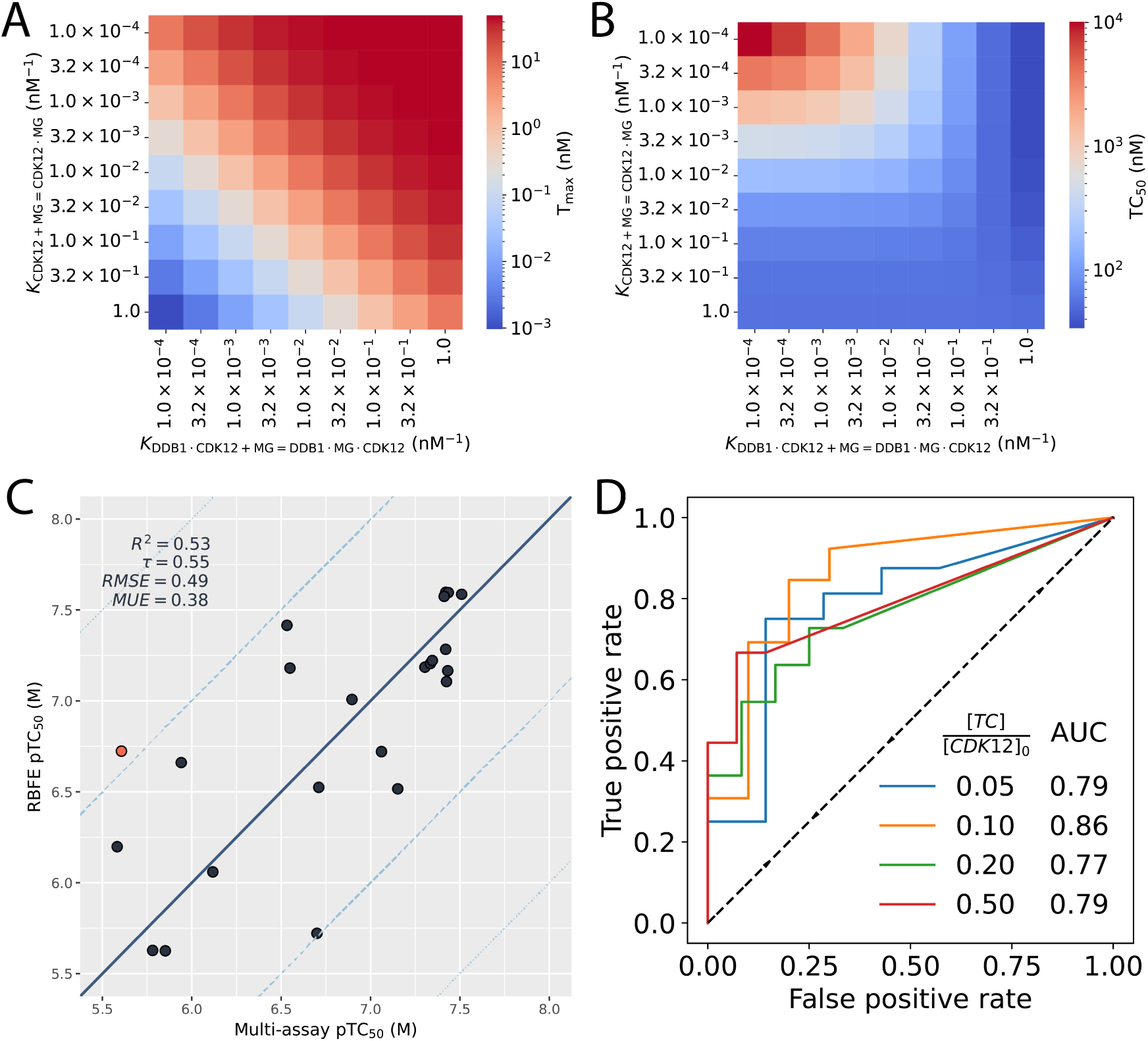
Predicting ternary complex formation from binding free energy calculations. (A) The maximum ternary complex concentration that a molecular glue can induce, given its binary and ternary binding constants. (B) The molecular glue concentration at which the ternary complex concentration reaches 50% of the maximum value at corresponding binary and ternary binding constants. (C) The molecular glue concentration at which the ternary complex concentration reaches 50% of the maximum value, derived based on the binary and ternary *K_D_*’s from experimental assays and from RBFE calculation. (D) The Receiver Operating Characteristic curves of classification of active molecular glues based on their concentration to induce a specified fraction of CDK12 in the ternary complex. A molecular glue is considered active if its pTC*_x_* > 7 (i.e. the molecular glue concentration required for the ternary complex concentration [TC] to reach x[CDK12]_0_ is less than 100 nM).

From the equilibrium constants of the reactions in Eq. 1, we can estimate both the maximum ternary complex concentrations Tmax and the molecular glue concentration TC_50_ for the ternary complex concentration to reach 50% of Tmax. We compared the TC_50_ values estimated from the binary and ternary *K_D_*values determined from experimental assays and from RBFE calculations, demonstrating reasonable agreement (Fig. 4c).

To assess the efficiency of classifying molecular glues in terms of ternary complex stabilities based on the predictions from RBFE calculations, we computed the Receiver Operating Characteristic (ROC) curves (Fig. 4d). We define TC*_x_* as the molecular glue concentration at which the ternary complex concentration reaches x[CDK12]_0_, and we consider a molecular glue active (in inducing ternary complex formation) if pTC*_x_* = − log_10_ TC*_x_* > 7 (corresponding to 100 nM TC*_x_*). The ROC curves for various values of *x* = [*TC*]/[*CDK*12] are shown, with area under curve (AUC) around 0.8, suggesting high classification accuracy.

#### Prediction of ubiquitination and degradation from DDB1-MG-CDK12 ternary complex ensembles

We showed that relative binding free energy calculations can predict ternary complex stability. Furthermore, experimental data reveal that ternary complex stability correlates well with degradation efficiency for the DDB1-MG-CDK12 system. Thus, estimating ternary complex stability computationally is a good proxy for predicting degradation efficiency. However, there are notable outliers. For instance, we compared two MGs, Roscovitine and DS50. Although Roscovitine and DS50 form ternary complexes with EC_50_ values of 0.7 µM and 1.9 µM, respectively, DS50 is an active degrader with a DC_50_ of 3.6 µM, while Roscovitine shows no degradation activity.^5^ Motivated by prior work finding a correlation between degradation potency and target protein accessibility to ubiquitin in a set of SMARCA2-VHL PROTACs,^29^ we hypothesized that this discrepancy arises from differences in the structural landscape of the ternary complexes induced by the two MGs. HREMD simulations of the ternary complexes induced by these MGs show that DS50 positions potential ubiquitination sites (lysine residues) closer to the distal end of the CRL complex (Fig. 5A), where ubiquitin would be loaded, thus explaining its superior degradation activity despite comparable ternary complex thermal stability.

**Figure 5:**
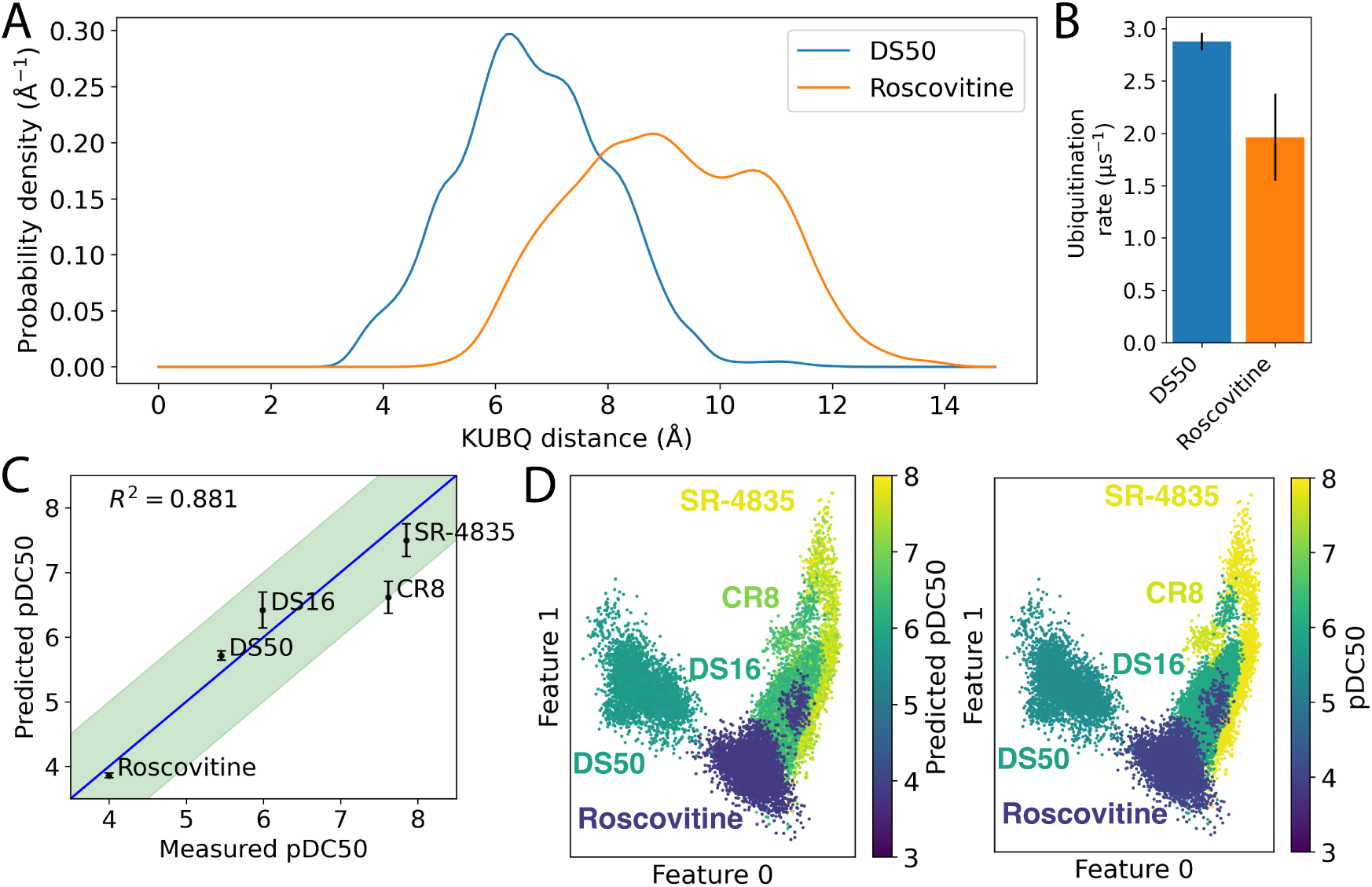
(A) Lysine-ubiquitin distance distributions for Roscovitine and DS50. The probability density shows that DS50 positions lysine residues closer to the C-terminal residue of Rbx1. (B) Estimated ubiquitination rates for Roscovitine and DS50. The ubiquitination rate represents the time-averaged frequency with which lysine residues remain within 10 Å of the C-terminal residue of Rbx1. (C) Results of the supervised VAE model. The model accurately predicts pDC_50_ for the test set of molecular glues excluded from training (R^2^ = 0.881). (D) The structured latent space of the model differentiates molecules based on their pDC_50_, with more potent MGs sampling higher values of Feature 0.

To evaluate how different MGs position potential ubiquitination sites (lysine residues) on CDK12-CycK within the ubiquitin-transfer assembly, we approximated the conformational landscape of the full cullin ring ligase (CRL) complex by superposing our DDB1–MG–CDK12-CycK simulations onto a simulation of the CRL scaffold. Specifically, we aligned snapshots from our ternary complex ensembles via DDB1 to a 100 ns trajectory of the full CRL complex induced by CR8. This superposition yields approximate CRL ensembles induced by each MG. Using the C-terminus of Rbx1 (located at the distal end of the CRL complex) as a proxy for ubiquitin to assess the ubiquitination potential of each MG, we found that DS50 positions the target protein more favorably for ubiquitination than Roscovitine (Fig. 5A).

To quantify this effect, we estimated the ubiquitination rate for each molecule, defined as the time-averaged frequency with which lysine residues remain within a predefined distance from the C-terminus of Rbx1 (see Methods). Using a distance cutoff of 10 Å, DS50 shows a substantially higher ubiquitination rate (∼2.9 µs^−1^) compared to Roscovitine (∼1.9 µs^−1^), consistent with DS50’s status as the more potent degrader (Fig. 5B).

Building on this observation, we developed a machine learning model to predict degradation potency based on the structural features of the ternary complex induced by a given MG. The model incorporates inter-chain distances and distances between putative ubiquitination sites and the C-terminus of Rbx1 at the distal end of the CRL complex (Fig. 8). Details of the model are provided in the Methods section.

Our model is a supervised variational autoencoder (VAE) that uses a randomly selected ensemble of frames from HREMD simulations of MG-induced ternary complexes to predict their pDC_50_. The model was trained on HREMD simulations of five MGs and tested on a separate set of five MGs not included in the training data. We designed the train-test split to be particularly challenging by placing pairs of MGs with similar EC_50_ values but divergent DC_50_ values (e.g., SR-4835 and DS16; DS50 and Roscovitine) together in the test set, assessing the model’s ability to distinguish degradation potency from binding affinity. The model accurately predicted the potency of the test set MGs, achieving an *R*^2^ value of 0.881 (Fig. 5C). Cross-validation on a further 20 train-test splits with seven training molecules and three test molecules yielded a median *R*^2^ = 0.767 ± 0.076.

A key benefit of the supervised VAE architecture is that the model learns a structured latent space, in which learned features correlate with degradation potency. As shown in Fig. 5D, embeddings from the test set simulations reveal that more potent MGs tend to occupy higher values of latent feature 0.

### Case Study: CRBN-dependent GSPT1 degradation

We applied GlueMap to identify CRBN-recruiting molecular glue (MG) degraders of GSPT1. Table 4 summarizes the main simulations performed in this section.

**Table 4:**
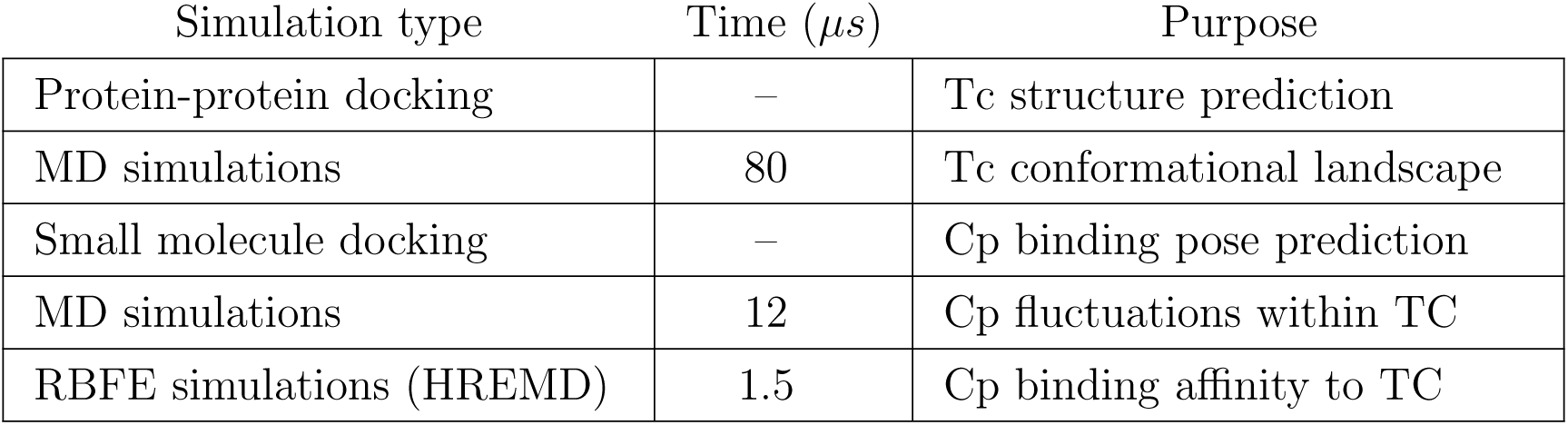
Summary of the molecular simulations performed for the CRBN-GSPT1 case study. The Time column indicates the cumulative duration of the simulations, whenever applicable. Tc = Ternary complex; Cp = Compound.

#### Prediction of ternary complex structure

We first validated GlueMap by modeling the ternary complex between CRBN, the known MG CC-885, and GSPT1. GlueMap can accurately predict the ternary complex structures of GSPT1-MG-CRBN, as demonstrated by the excellent agreement with the crystal structure (PDB ID 5HXB^53^) of the ternary complex formed by GSPT1, CC-885, and CRBN (Fig. S2).

#### Computation of CRBN-GSPT1 free energy landscape

We performed large scale unbiased MD simulations starting from docked poses of CRBN-GSPT1. We used tICA^54^ to featurize the conformational free energy landscape and identified diverse PPI states of the protein complex using K-means clustering (Fig. 6A). The 5HXB crystal structure was the most populated PPI state (Fig. 6B).

**Figure 6:**
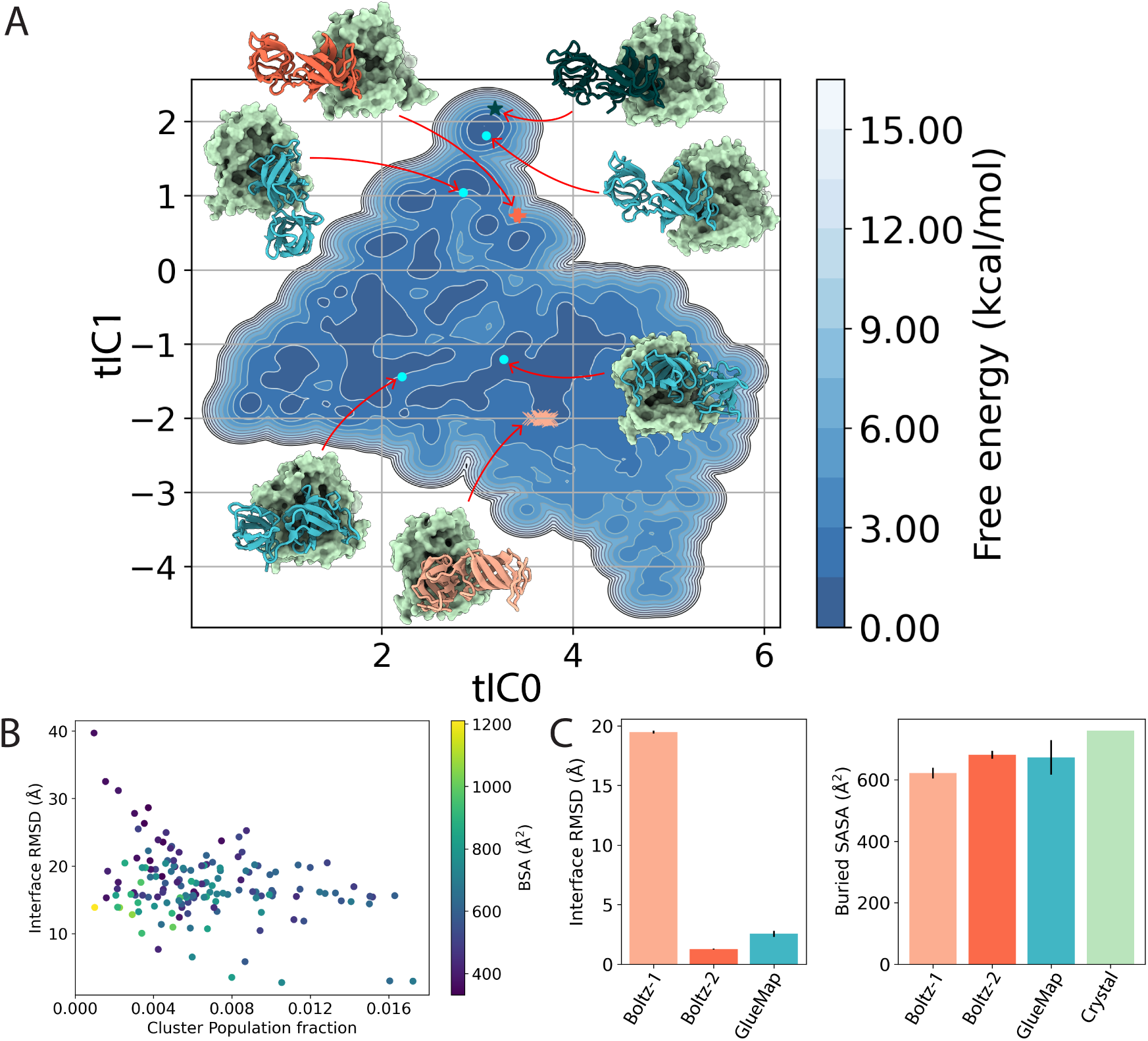
(A) GlueMap free-energy landscape of the GSPT1–CRBN encounter complex with inset representative structures from MD, Boltz-1 and Boltz-2 predictions, and the GSPT1-CC885-CRBN ternary complex crystal structure (PDB ID 5HXB). CRBN is shown as a green surface; GSPT1 is cyan for MD snapshots, pale orange for Boltz-1, red for Boltz-2, and dark green for 5HXB. The projected locations of the Boltz-1/-2 predictions on the landscape are marked by orange “×” and red “+”, respectively. (B) Cluster population versus mean interface RMSD for PPI clusters. (C) Comparison of interface RMSD and buried solvent-accessible surface area of Boltz-1/-2 predictions, frames from the most-populated PPI state, and 5HXB.

We compared the structures obtained from our simulations to predictions from AI cofolding models Boltz-1 and Boltz-2. Our simulations recovered structures more similar to the published crystal structure than Boltz-1 (Fig. 6C); while Boltz-2 was able to recapitulate the crystal structure, this structure was included in the training data of that version of Boltz-2.^13^

#### Recovery of known molecular glues using GlueMap

To validate GlueMap’s capabilities in identifying known molecular glues, we retrospectively screened a virtual library of cereblon binders comprising approximately 4,500 compounds sourced from Enamine. This library was augmented with three known molecular glues: CC-90009, CC-885, and MRT-2359. Using the crystal structure 6XK9 as the basis, we followed the GlueMap workflow outlined in Fig. 1.

Initial filtering based on docking scores and predicted interactions with GSPT1 narrowed the library to 100 candidates. These candidates were further refined using long molecular dynamics (MD) simulations, with stability assessed via ligand root-mean-square fluctuation (RMSF). Of the 100 compounds, 24 exhibited high stability and were subjected to relative binding free energy (RBFE) calculations. Among the screened compounds, the three known molecular glues—CC-90009, CC-885, and MRT-2359—ranked among the top-performing molecules, as shown in Fig. S3.

These results underscore GlueMap’s ability to reliably recover previously identified molecular glues by integrating docking, MD simulation stability, and RBFE predictions, thereby offering a robust and comprehensive evaluation framework. This methodology extends prior work^55^ by incorporating thermodynamic and dynamic stability assessments alongside traditional docking metrics.

#### Discovery of CRBN-GSPT1 molecular glues by hierarchical virtual screening

We recently discovered a CRBN-GSPT1 molecular glue, indicated as MG1 in Fig. 7A. By Western blotting, we confirmed that MG1 degrades GSPT1 in HEL92.1.7 cells. Additionally, we confirmed its proteasome-dependent mechanism of action via the Cullin–RING ligase pathway, as evidenced by the absence of GSPT1 degradation, when cells were co-treated with MLN4924 and Bortezomib (Fig. 7B). Additional experimental validation in HEK293-GSPT1-HiBIT cells confirmed MG1 as a potent glue degrader with a DC_5_0 of 19.3 nM (average of n=2, Table S1) and a Dmax of 100% (Fig. 7C). To test the hypothesis that MG1 would bind at the known CRBN-GSPT1 interface,^56^ we docked MG1 to receptor structure 6XK9 using MOE,^46^ which resulted in a strongly bound pose with a docking score S = −12.05. This ternary-complex model is provided in the Supplementary Information. As shown in Fig. 7D, MG1 adopts an orientation similar to the known glue degrader CC-90009.^56^ While the glutarimide moiety forms hydrogen bonds with CRBN residues Trp380 and His378, the urea group is positioned at the protein-protein interface, where it forms two additional hydrogen bonds with CRBN His353 and GSPT1 Gln533.

**Figure 7:**
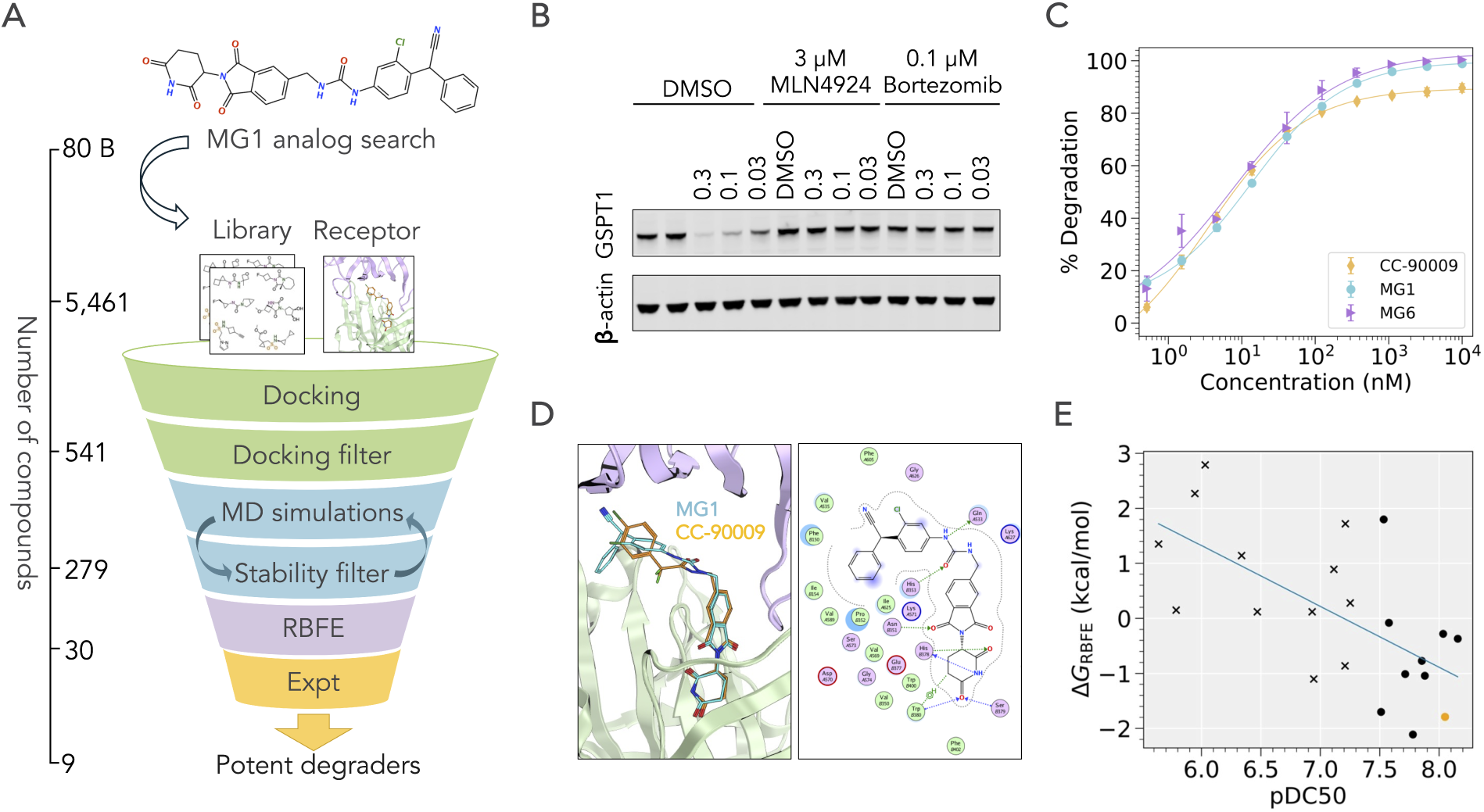
Prospective CRBN-GSPT1 virtual screening. (A) Compound MG1 is used as a query to build a focused library, which is subsequently docked to the receptor. The following steps of MD simulation cycles and RBFE calculations identify stable ternary complexes with high-affinity binders. Experimental validation results in nine potent degraders with DC_50_ < 50 nM. (B) Western blot analysis of MG1. β-actin is used as a loading control to confirm normal protein expression in cells. Concentrations are reported in µM. (C) Dose–response curves showing GSPT1 degradation induced by CC-90009, MG1 and MG6, the most potent compound identified in silico (average of n=2 replicates). Lines correspond to Hill equation fits. (D) Left panel: MG1 (cyan) docked to GSPT1 (violet) and CRBN (green). Compound CC-90009 (orange) is shown for comparison. Right panel: MG1 interaction profile. (E) The correlation between degradation potency pDC_50_ (average of n=2 replicates) and the binding affinity Δ*G_RBFE_* of compounds to CRBN-GSPT1 is *R*^2^ = 0.46. Circles and crosses indicate, respectively, potent and weak degraders (threshold pDC_50_ = 7.3). Compound CC-90009 (orange circle) is shown for comparison.

Taken together, the above results suggested that MG1 was suitable to seed a virtual screening campaign. Accordingly, we set up a pipeline combining docking and cycles of MD simulations, followed by RBFE calculations and experimental validation (Fig. 7A). As the first step of this campaign, we generated a focused library of compounds by searching the Enamine REAL and WuXi GalaXi collections^57^ for analogs of MG1. A total of 5,461 glutarimide-containing compounds were selected and docked to CRBN–GSPT1 (PDB ID 6XK9) using pharmacophore restraints on the glutarimide moiety.^46^ The resulting ternary complexes were ranked with a scoring function that combines docking score and number of specific ligand–protein contacts. Approximately 10% of these complexes achieved a total score > 1 (Eq. 10) and were consequently evaluated for ligand stability using short MD simulation cycles. Compounds exhibiting large structural fluctuations (see Methods) were excluded, yielding 279 stable ternary complexes.

To reduce the cost of RBFE simulations, we first downsampled the stable ternary complexes using the divide-and-conquer strategy described in the Methods section. We then ran 58 RBFE simulations and analyzed the results using DiffNet,^51^ in order to compute the per-compound binding affinities (Δ*G_RBFE_*). A network representation of the RBFE calculations, together with the ΔΔ*G_RBFE_* and Δ*G_RBFE_* values are provided in the Supporting Information. As shown in Fig. S4, the DiffNet-derived ΔΔG values are in excellent agreement with ΔΔ*G_RBFE_* (R^2^ = 0.94), supporting the reliability of the Δ*G_RBFE_*estimates and their use in Fig. 7E for correlating compound binding affinity with degradation potency.

The final step of our virtual screening campaign involved experimental testing of thirty compounds using HiBiT. Twenty two were active, with DC_50_ values ranging from single-digit nanomolar to micromolar, whereas eight were inactive, having DC_50_ `≥` 10µM (see summary Table S1). The inactive compounds showed a higher average Δ*G_RBFE_* = 1.70 kcal/mol compared to 0.12 kcal/mol of the active compounds, a difference that is statistically significant (p = 0.03, Fig. S5) and supports the conclusion that our virtual screening campaign successfully distinguished active from inactive molecular glues. Importantly, as shown in Fig. 7E, the relationship between measured GSPT1 degradation and binding affinity indicates that stronger binders (lower Δ*G_RBFE_*) induce more potent degradation. However, the moderate correlation coefficient (R^2^ = 0.46) suggests that additional factors likely contribute to this protein degradation. Specifically, GSPT1 engages CRBN via the degron motif located in one of its two domains, while the other domain is distant from both the molecular glue and CRBN surface, suggesting that ternary complex stability and GSPT1 intrinsic flexibility may modulate the observed degradation potency.

In conclusion, this prospective virtual screening campaign demonstrates GlueMap’s ability to predict novel molecular glues through a combination of docking, dynamic stability analysis, and binding free energy calculations.

## Discussion

GlueMap is a computational platform for modeling the structural ensembles of ternary complexes, predicting their thermodynamic stability, and evaluating downstream effects such as ubiquitination and degradation efficiency of molecular glues (MGs). By integrating structural and thermodynamic metrics with ubiquitination potential, GlueMap enables a multi-criteria approach to rational MG design.

This study, together with prior work,^29,30^ demonstrates that computational modeling can predict the conformational landscapes of ternary complexes with high atomistic accuracy. These landscapes are instrumental for identifying molecular glues that stabilize specific protein–protein interactions with desirable properties, including enhanced ubiquitination potential and favorable thermodynamic stability.

The case studies of DDB1–CDK12 and CRBN–GSPT1 illustrate GlueMap’s utility across distinct degradation systems. In the DDB1–CDK12 system, GlueMap revealed strong correlations between ternary-complex stability and degradation efficiency, as well as between predicted binding free energies and measured complex stability. These relationships underscore GlueMap’s ability to capture the thermodynamic determinants of molecular glue activity. Importantly, GlueMap accurately recovered ternary-complex structures for multiple CDK12 degraders, whereas Boltz-1 and Boltz-2 failed to reproduce the CDK12–DDB1 crystal structure and sampled a reduced diversity of protein–protein interface geometries. The disparity between Roscovitine and DS50 degradation efficiencies, despite similar thermal stabilities, highlights the importance of structural features beyond overall complex stability. GlueMap simulations showed that DS50 positions lysine residues closer to the ubiquitin-loading site relative to Roscovitine, explaining its superior degradation activity. This capacity to resolve subtle structural differences that govern ubiquitination efficiency distinguishes GlueMap from approaches that rely solely on static structure prediction or binding affinity.

Machine learning further enhances GlueMap’s predictive capabilities. A supervised variational autoencoder (VAE), trained on molecular dynamics trajectories incorporating both inter-chain distances and lysine–ubiquitin proximities, achieved high accuracy (R^2^ = 0.881 on the test set) in predicting degradation potency from structural features alone. Cross-validation across 20 train–test splits yielded a median *R*^2^ = 0.767 ± 0.076, demonstrating robust generalization. Notably, the model distinguished degradation potency from binding affinity by learning a structured latent space in which potent degraders occupy distinct regions characterized by favorable ubiquitination geometries.

For CRBN–GSPT1, GlueMap successfully recovered known MG degraders and identified nine novel candidates through prospective virtual screening using a simulated ensemble of protein–protein interaction states. All nine compounds exhibited potent degradation activity with DC_50_ values below 50 nM, validating the predictive power of our conformational ensemble approach. This campaign succeeded through GlueMap’s integrated workflow: protein–protein docking for initial ternary-complex generation, a scoring function combining docking scores and protein–ligand contact analysis, molecular dynamics simulations to eliminate unstable complexes, and RBFE calculations to prioritize high-affinity binders. The moderate correlation between RBFE values and measured degradation potency (R^2^ = 0.46) indicates that binding affinity is not the sole determinant of GSPT1 degradation. Additional factors, including intrinsic flexibility and lysine orientation relative to the E3 ligase active site, likely contribute to degradation efficiency. The alternative CRBN–GSPT1 interfaces revealed by our encounter-complex simulations (Fig. 6) present opportunities to probe these mechanisms experimentally and to further optimize degraders by selecting for geometrically favorable PPIs.

Future developments could extend GlueMap to additional protein targets and a broader range of E3 ligases, including membrane-associated systems. Incorporating experimental feedback loops and applying active-learning strategies may further enhance predictive accuracy. Integration with emerging structure-prediction methods, when validated against conformational ensemble data, could accelerate initial complex generation while preserving GlueMap’s strengths in thermodynamic and dynamic modeling.

## Methods

### DDB1–CDK12 case study

#### DDB1–MG–CDK12–CycK complex structure prediction

Initial poses were generated using Method 4B in MOE.^38^ Poses with the highest double-cluster populations were selected for HREMD simulations. Each system was prepared and energy-minimized in MOE, solvated in a cubic box with a 10 Å buffer, and neutralized with Na^+^ and Cl^−^ ions. Simulations were run for 30 ns per system with 32 replicas spanning 298.15–400 K logarithmically and using a 2 fs time step. Proteins were parameterized using the Amber ff14SB force field,^58^ and small molecules were parameterized using proprietary software; parameter files are provided in the Supplementary Information.

Inter-chain residue distances were computed between DDB1 and CDK12. Pairs of residues whose Cα atoms were within 10 Å in any frame were designated as interface residues. Cα distances between all interface pairs were collated across all frames and subjected to principal component analysis (PCA), retaining the top six components. The resulting low-dimensional ensemble was clustered by k-means with k = 50, and the highest-population cluster centroid was selected as the predicted ternary-complex model.

#### Structure predictions using Boltz-1 and Boltz-2

DDB1–MG–CDK12 and CRBN–MG–GSPT1 ternary complex structures were predicted by generating 20 independent samples using Boltz-1 or Boltz-2 with default parameters. Cyclin K was omitted from the DDB1–MG–CDK12 predictions due to the large memory demands associated with predicting the full quaternary structure; since evaluation was restricted to the DDB1–CDK12 interface, this omission is not expected to affect ligand or interface RMSD values.

### Cullin–RING ligase simulations

We performed 100 ns molecular dynamics simulations of the Cullin–RING ligase complex comprising the molecular glue CR8, Cyclin K, CDK12, DDB1, Cul4, and RBX1. Simulations were run in Amber with a 2 fs time step. The system was prepared in MOE by superposing crystal structure 6TD3 onto cryo-EM structure 7OKQ via DDB1.^5,59^ DDB1 and DCAF1 were deleted from the 7OKQ system. The system was solvated with a 10 Å buffer and neutralized with Na^+^ and Cl^−^ ions. This simulation served as a reference for determining Lys–UBQ distances and ubiquitination rates.

#### Supervised representation learning

HREMD simulations^29,60^ were performed in Amber for ten crystal structures of the DDB1– CDK12–Cyclin K complex (PDB IDs: 6TD3, 8BU1, 8BU2, 8BU5, 8BU9, 8BUD, 8BUG, 8BUN, 8BUR, and 8BUT^5^). Each system was prepared and energy-minimized in MOE, solvated with a 10 Å water buffer, and neutralized with Na^+^ and Cl^−^ ions. Simulations were run for 50 ns per system with 32 replicas spanning 298.15–400 K logarithmically, using a 2 fs time step. Proteins were parameterized with Amber ff14SB,^58^ and small molecules were parameterized using proprietary software (parameters provided in the Supplementary Information).

Lysine–ubiquitin distances were computed by superposing ternary-complex (TC) frames onto a reference MD simulation of the CRL complex. Each TC frame was aligned to ten CRL poses using rigid-body superposition on DDB1. For each aligned pair, we measured the distance from the side-chain Nζ of each CDK12 and CycK lysine to the backbone carbonyl oxygen of RBX1 Val39 (used as a ubiquitin proxy), yielding 29 Lys–UBQ distances.

Using these simulations, we trained a supervised variational autoencoder (VAE) to predict molecular-glue pDC_50_ values from the free-energy landscape of DDB1–glue–CDK12–Cyclin K complexes. Input features consisted of (i) inter-chain Cα distances between DDB1 and CDK12, where residue pairs were classified as interface contacts if their Cα atoms approached within 10 Å in any TC frame, and (ii) per-lysine Lys–UBQ distances averaged over the CRL-aligned poses (45 total sites from CDK12 and CycK). For each snapshot, these features were concatenated into a single vector representing the ternary-conformation in the CRL context.

The VAE latent representation was passed to an Ensemble Transformer regression head that predicted pDC_50_ values based on an ensemble of n_samp_ = 128 frames using a self-attention layer, mean pooling, and three fully connected layers (Fig. 8). Because the regression module aggregates an arbitrary number of frames, inference can be performed with any n_samp_ subject to GPU memory limits. Unless otherwise noted, we used n_samp_ = 128; smaller values increase prediction variance, whereas larger values improve stability at higher computational cost.

**Figure 8:**
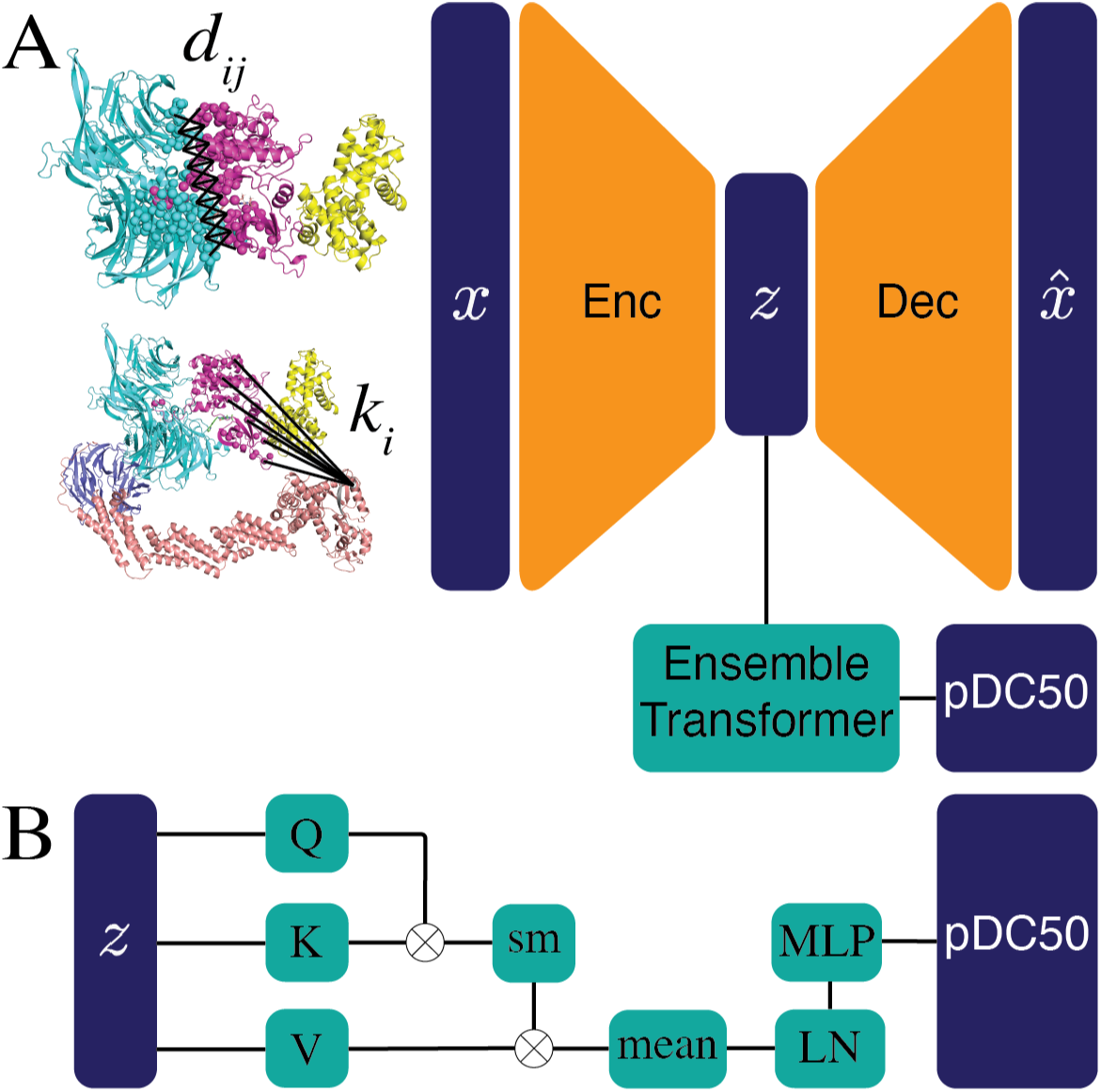
(A) Schematic of the supervised VAE. Simulation frames are featurized using interchain residue distances (d*_ij_*) and lysine–ubiquitin distances (k*_i_*). (B) Ensemble Transformer regression module for pDC_50_ prediction using self-attention. Abbreviations: SM (softmax), LN (layer normalization), MLP (multi-layer perceptron).

### Model training

The model was first trained using a 50:50 train–test split in which DS18, DS17, HQ461, DS61, and Z7 were assigned to the training set, and CR8, SR-4835, DS16, Roscovitine, and DS50 were assigned to the test set. To assess robustness to data partitioning, we additionally trained on 20 independent 70:30 train–test splits, randomly generated with the constraint that each test set contain one non-degrader (no detectable degradation), one weak degrader (DC_50_ > 400 nM), and one potent degrader (DC_50_ < 400 nM). Molecular glues lacking experimentally detected DC_50_ values were labeled as pDC_50_ = 4 (corresponding to 10 µM) for training.

Training was performed for 50 epochs with an 8-dimensional latent space. The encoder and decoder each comprised two fully connected layers (4096 units). The regression module consisted of one self-attention layer followed by three fully connected layers (4096 units).

#### Estimation of ubiquitination rates

We estimated a relative ubiquitination rate for each molecular glue (MG) using HREMD simulations of the DDB1–MG–CDK12 ternary complex. Each HREMD snapshot was rigidly aligned to a set of reference CRL poses via DDB1, embedding the ternary structure within the full CRL context. For every aligned snapshot pair, we computed the distance between the side-chain Nζ atom of each CDK12 and CycK lysine and the backbone carbonyl oxygen of RBX1 Val39, counting a “ubiquitination event” whenever this distance fell below a cutoff d_cut_ = 10 Å.

The relative ubiquitination rate was defined as

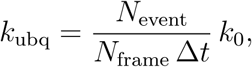

where N_event_ is the total number of ubiquitination events, N_frame_ is the number of analyzed frames, Δt = 0.1 ns is the time interval between frames, and k_0_ = 0.05 is a numerical scaling factor. Each lysine within the cutoff distance contributes one event. Because HREMD accelerates conformational sampling, k_ubq_ is a comparative metric—not an absolute rate— used solely to rank MGs under a consistent simulation protocol.

#### Relative Binding Free Energy calculations

RBFE calculations consider the following binding reactions and alchemical transformations:

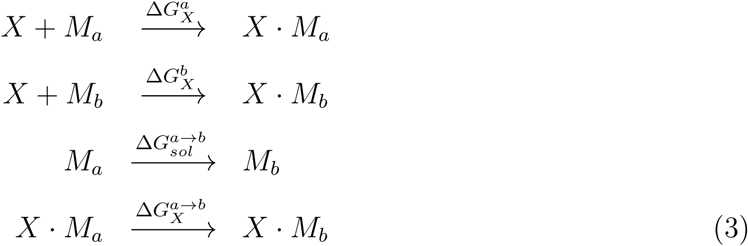

where X ∈ {P, E, P · E} denotes the receptor, and M*_a_* and M*_b_* are the two glue molecules being compared. The relative binding free energies are given by

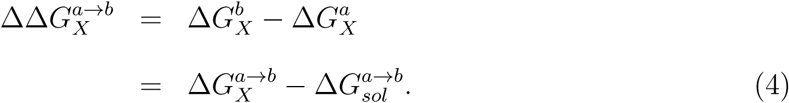

There are two binary RBFEs, 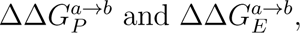 corresponding to receptors P and E. Our workflow enables computation of either or both.

To evaluate the thermodynamic stability of ternary and binary complexes involving molecular glues, we employed a workflow for calculating relative binding free energies (RBFEs).^50–52^ This approach is based on the thermodynamic cycles in Eqs. 3 and 4 and integrates molecular dynamics (MD) simulations with modern free-energy analysis techniques. The workflow is fully automated, spanning molecular design, simulation setup, RBFE estimation, and visualization.

#### RBFE workflow overview

The overall RBFE workflow is summarized in Fig. S6 and consists of the following steps:

1. **Docking and structure preparation:** Designed molecules are docked into binary and ternary complexes using a common-core docking strategy that preserves conserved substructures while allowing substituent flexibility. Experimental or computationally derived ternary structures serve as docking templates.
2. **Force-field parameterization:** Parameters for molecular-glue candidates are generated on-the-fly using a proprietary force-field generator, ensuring compatibility with the Amber14 protein force field.
3. **Atom matching and morph-graph construction:** Atom mappings between molecules create a morph graph whose nodes represent compounds and whose edges specify al-chemical transformations. The graph is optimized to maximize structural similarity while maintaining connectivity.
4. **System preparation:** For each graph edge, three alchemical systems are built: molecule in water, molecule in the binary complex, and molecule in the ternary complex. Binary complexes are generated by removing one receptor from the ternary structure. Systems are solvated with explicit water and ions to 0.15 M NaCl.
5. **Simulation protocol:** Systems undergo energy minimization and relaxation MD, followed by HREMD simulations optionally using REST2^61^ to improve conformational sampling.
6. **Free-energy calculations:** Free-energy differences are computed using MBAR or BAR. RBFE results are aggregated and analyzed using DiffNet to obtain maximum-likelihood RBFEs for binary and ternary targets.
7. **Results and reporting:** The computed RBFEs are summarized in an Excel report containing numerical values and structural analyses for compound prioritization.

This workflow combines enhanced sampling, optimized atom mappings, and scalable automation to deliver high-accuracy RBFEs and mechanistic insight into molecular glue activity.

#### Derivation of binary and ternary binding constants from experimental assays

We determined equilibrium binding constants by jointly fitting binary binding assays and ternary-complex formation assays for CDK12 and DDB1 (Fig. S1). Glue-mediated ternary-complex formation is modeled using the reactions in Eq. 1.

The concentrations of all molecular species are obtained by solving

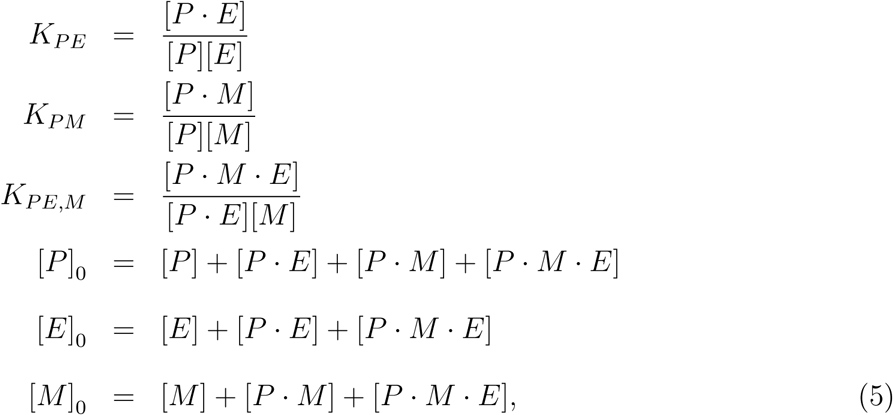

where [X]_0_ is the initial concentration of X ∈ {P, E, M}.

In binary and ternary binding assays, [P]_0_ and [E]_0_ are fixed, and [M]_0_ is titrated; equilibrium concentrations therefore depend solely on [M]_0_.

In the TR-FRET assay, the signal is modeled as

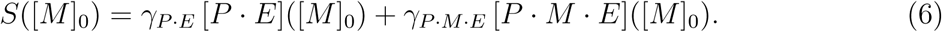

Binary binding between P and M is described by the competitive assay:

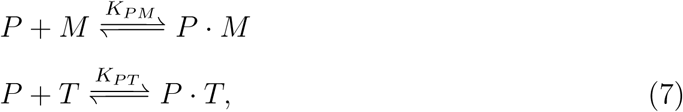

with signal

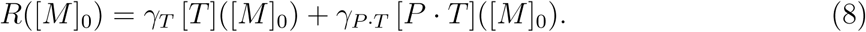

Given dose–response curves from TR-FRET and tracer-competition assays, we fit *K_PE_*, *K_PM_*, *K_PE,M_*, *K_PM,E_*, *K_PT_* and signal-to-concentration ratios γ*_P_*_·*E*_, γ*_P_*_·*M*·*E*_, γ*_T_*, γ*_P_*_·*T*_. The objective function is

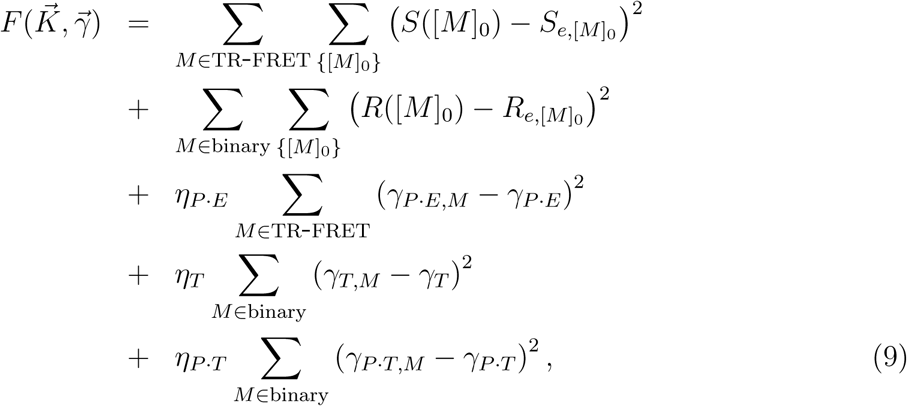

subject to the thermodynamic cycle-closure constraint in Eq. 2.

In Eq. 9, S*_e,_*_[*M*]0_ and R*_e,_*_[*M*]0_ are the experimental readouts at concentration [M]_0_. Derived binary *K_D,b_* and ternary *K_D,t_* values are summarized in the Supplementary Information.

### CRBN–GSPT1 case study

#### Molecular dynamics simulations

Forty structures generated from protein–protein docking of GSPT1 and CRBN were used as starting points for molecular dynamics (MD) simulations. Each structure was prepared in MOE.^46^ To reduce computational cost, DDB1 was removed.

For each starting structure, two replicas were simulated for 2 µs each, totaling 80 simulations. MD simulations were carried out in OpenMM with hydrogen mass repartitioning and a 4 fs time step. Harmonic restraints maintained the structural integrity of the Zn ion in CRBN. Each system was solvated in a rectangular water box with a 9 Å buffer, neutralized, and supplemented with 100 mM NaCl, yielding systems containing approximately 170,000 atoms.

Proteins were parameterized with the Amber14 force field; parameters for CC-885 were generated using internal and commercial tools. Small-molecule parameter files are provided in the Supplementary Information.

#### Encounter-complex clustering

Encounter complexes from CRBN–GSPT1 MD simulations were analyzed using time-lagged independent component analysis (tICA).^54^ Input features comprised the inter-chain distance matrix describing the CRBN–GSPT1 interface and the interface RMSD relative to the crystal structure 5HXB. Interface residues were defined as those whose Cα atoms came within 15 Å of one another in any MD frame.

tICA reduced the RMSD and distance-matrix features to an eight-dimensional latent space. Features were scaled using the kinetic-map method,^62^ and frames were clustered into 150 states using k-means as implemented in scikit-learn.^63^

#### Compound library for virtual screening

We used InfiniSee^57^ to search for analogs of a novel GSPT1 degrader (Fig. 7) within the Enamine REAL and WuXi GalaXi libraries, using ECFP4 fingerprints with a minimum similarity threshold of 0.5. The resulting compounds were processed in two steps:

1. Using MOE’s sdwash,^46^ salts and small components were removed, protomers at pH 7 were enumerated, and hydrogens were added.
2. Using MOE’s sdfilter,^46^ protomers with predicted frequencies greater than 20% were retained.

#### Prospective virtual screening workflow and analysis

The prospective virtual screening workflow consists of four stages:

1. **Pharmacophore-guided docking:** The receptor 6XK9 was prepared using the MOE QuickPrep protocol.^46^ The compound library (prepared as described above) was rigidly docked in MOE, guided by a three-feature pharmacophore derived from co-crystal structures 5HXB and 6XK9: two hydrogen-bond acceptor sites corresponding to interactions between the glutarimide of CC-885 or CC-90009 and CRBN residues His378 and Trp380, and one hydrogen-bond donor site corresponding to the interaction of CC-90009 with His378 (Fig. 7).
2. **Scoring and filtering of docked poses:** Docked poses were filtered using a scoring function combining the MOE docking score S and the number of specific ligand–protein contacts N*_c_*, implemented via softplus functions:

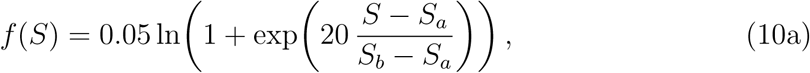

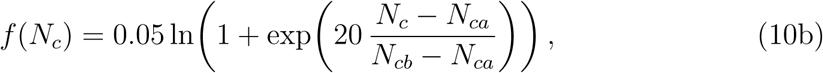

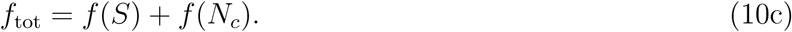 Complexes with f_tot_ `≥` 1 were retained. Parameters were tuned to S*_a_*= −4, S*_b_* = −10, N*_ca_* = 1, and N*_cb_*= 3. Specific interactions contributing to N*_c_* were computed using ProLIF,^64^ including HBDonor, HBAcceptor, Anionic, Cationic, CationPi, PiCation, PiStacking, XBAcceptor, and XBDonor.
3. **MD simulations to assess ternary stability:** Filtered complexes were solvated in a rectangular box with a 12 Å buffer and 150 mM NaCl. Heavy atoms were restrained during heating. Ten cycles of 2 ns simulations were performed in the NPT ensemble using Amber’s pmemd.cuda with hydrogen mass repartitioning and a 4 fs time step. At the end of each cycle, we computed the fraction of time the ligand RMSD, ⟨f_RMSD_⟩, remained below 2.5 Å relative to the minimized pose. Complexes with ⟨f_RMSD_⟩ < 0.5 were removed. Remaining complexes were ranked by: (1) low mean ligand RMSD; (2) low ligand RMSD standard deviation; (3) high mean CRBN–GSPT1 contacts; (4) high mean ligand–protein contacts. The top 15 per category were retained. CRBN–GSPT1 contacts were computed using a 4.5 Å cutoff.
4. **RBFE simulations:** RBFEs were computed following the workflow in Section “Relative Binding Free Energy calculations”. HREMD simulations were 25 ns per replica. Final ΔΔ*G*_RBFE_ and DiffNet-transformed Δ*G*_RBFE_ values for experimentally tested compounds are provided in the Supplementary Information. These Δ*G*_RBFE_ values are shifted and should not be interpreted as absolute binding free energies.

#### Western blotting

HEL92.1.7 (ATCC, TIB-180) cells were pre-treated with 3 µM MLN4924 (Med Chem Express, HY-70062) or 0.1 µM Bortezomib (Med Chem Express, HY-10227) for 30 min. Cells were then treated with the indicated Atommap compounds for 16 h. Protein levels of GSPT1 (Proteintech, 10763) and Actin (Cell Signaling Technologies, 8H10D10) were quantified by western blotting using a LI-COR imaging system.

#### Degradation assay

HEK293 GSPT1-HiBIT cells (C-terminally tagged) were generated by CRISPR knock-in, and a monoclonal line was isolated for all assays. Cells were incubated with compounds for 24 h, after which HiBIT signal was measured using the Nano-Glo HiBIT Lytic Detection Reagent (Promega, N3030) according to the manufacturer’s instructions.

#### Experimental data analysis and comparison with binding free energy predictions

Measured degradation potencies were expressed as DC_50_ values and transformed to pDC_50_ via

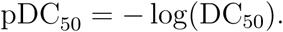

When multiple DC_50_ values were available, we used

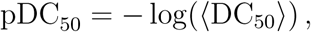

where ⟨DC_50_⟩ denotes the mean of replicate measurements.

To correlate experimental degradation potencies with RBFE predictions (e.g., Fig. 7E), pairwise ΔΔ*G*_RBFE_ values were transformed into per-compound Δ*G*_RBFE_ values using DiffNet.

Two-sample t-tests were performed in Python using the scipy library. A p-value < 0.05 was considered statistically significant.

## Supporting information

Ternary complex model of MG1 for GSPT1-CRBN

Excel spreadsheet for prospective virtual screening of GSPT1-CRBN molecular glues

Excel spreadsheet of the binary and ternary KD values fit to experimental data, for comparison to binding free energy calculations

Small molecule parameters for GSPT1-CRBN molecular glues

Small molecule parameters for CDK12-DDB1 molecular glues

## Author contributions

J.A.I., Y.W., and H.X. designed the study, supervised the work, and reviewed the manuscript. H.X. fitted the experimental data, and Y.W. performed the RBFE calculations for CDK12–DDB1 stability and supervised all RBFE analyses. T.P. developed software tools and implemented their application to the systems described. C.K. conducted the experimental studies. A.M.R. performed encounter-complex simulations for GSPT1–CRBN and contributed to the development of the GlueMap framework. J.A.I. carried out ternary-complex predictions and retrospective validation for GSPT1–CRBN. Z.M. performed most of the CDK12–DDB1 analyses and analyzed encounter-complex simulations for GSPT1–CRBN. F.T. conducted prospective virtual screening for GSPT1–CRBN. J.A.I., Z.M., and F.T. wrote most of the manuscript. All authors reviewed the final version. J.A.I., Y.W., and H.X. are corresponding authors.

## Acknowledgement

We thank Lakshmi Miller-Vedam for generating the visualizations of the GSPT1–CRBN and CDK12–DDB1 complexes, and Arie Zask for overseeing the purchase of GSPT1 glues from Enamine.

## Supplementary Information

The Supplementary Information includes: seven figures and one table; an Excel spreadsheet containing the binary (K*_D,b_*) and ternary (K*_D,t_*) dissociation constants fitted to experimental data for comparison with the binding free energy calculations (DDB1–Cyclin K case study); zip files containing the small-molecule parameters used in the CDK12–DDB1 and GSPT1–CRBN simulations; the docking model of the CRBN–MG1–GSPT1 ternary complex; and an Excel file summarizing the prospective CRBN–GSPT1 virtual screening results.

