## Supplementary Information for "Discovery of molecular glues by modeling ternary complex conformational ensembles and thermodynamic stability"

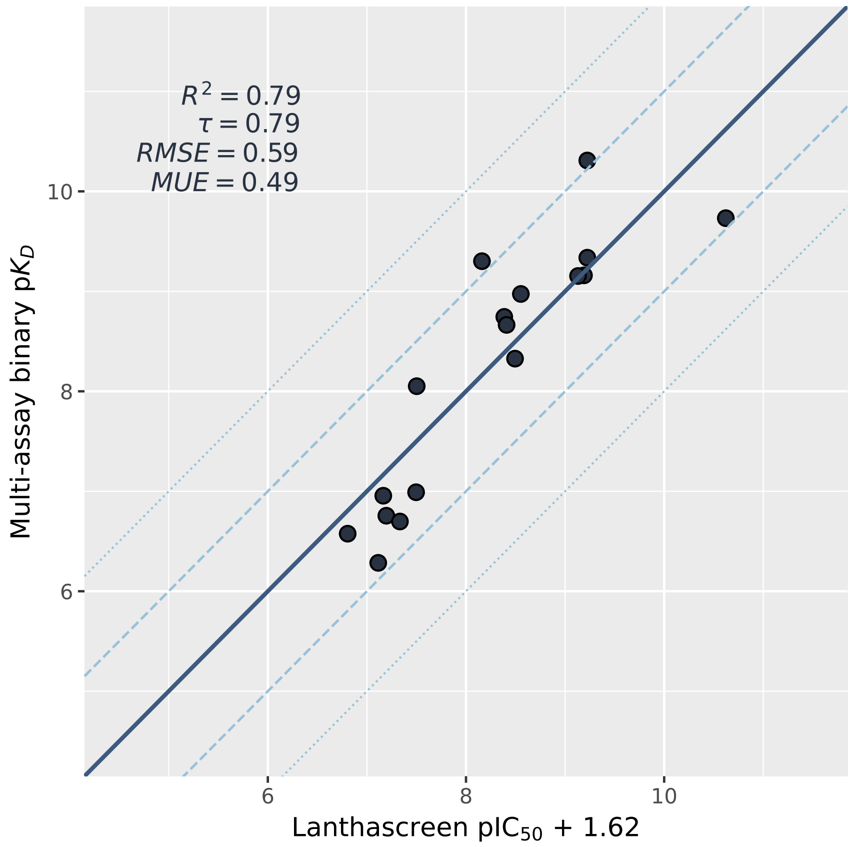


Figure S1: Comparing the binary $K_{\mathrm{D,b}}$ and ternary $K_{\mathrm{D,t}}$ values determined by the global fit of chemical reaction models to the literature values of $IC_{50}$ and $EC_{50}$ determined from phenomenological models.

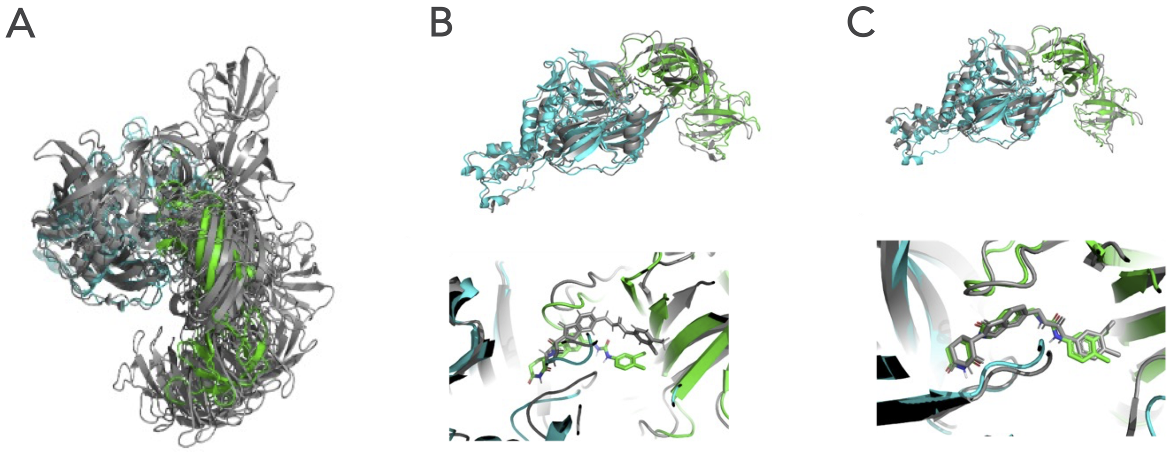


Figure S2: Accurate prediction of the GSPT1 – CC-885 – CRBN ternary complex using GlueMap. (A) Protein-protein poses derived from docking GSPT1 and CRBN. (B) Initial ternary complex model of GSPT1 – CC-885 – CRBN after compound docking and scoring. I-RMSD = 8.0 Å; ligand RMSD = 2.1 Å. (C) Refined ternary complex model after long MD simulations (100 ns x 2 replicas). I-RMSD = 3.5 Å; ligand RMSD = 1.0 Å. GSPT1, CRBN and CC-885 are colored in green and gray, respectively.

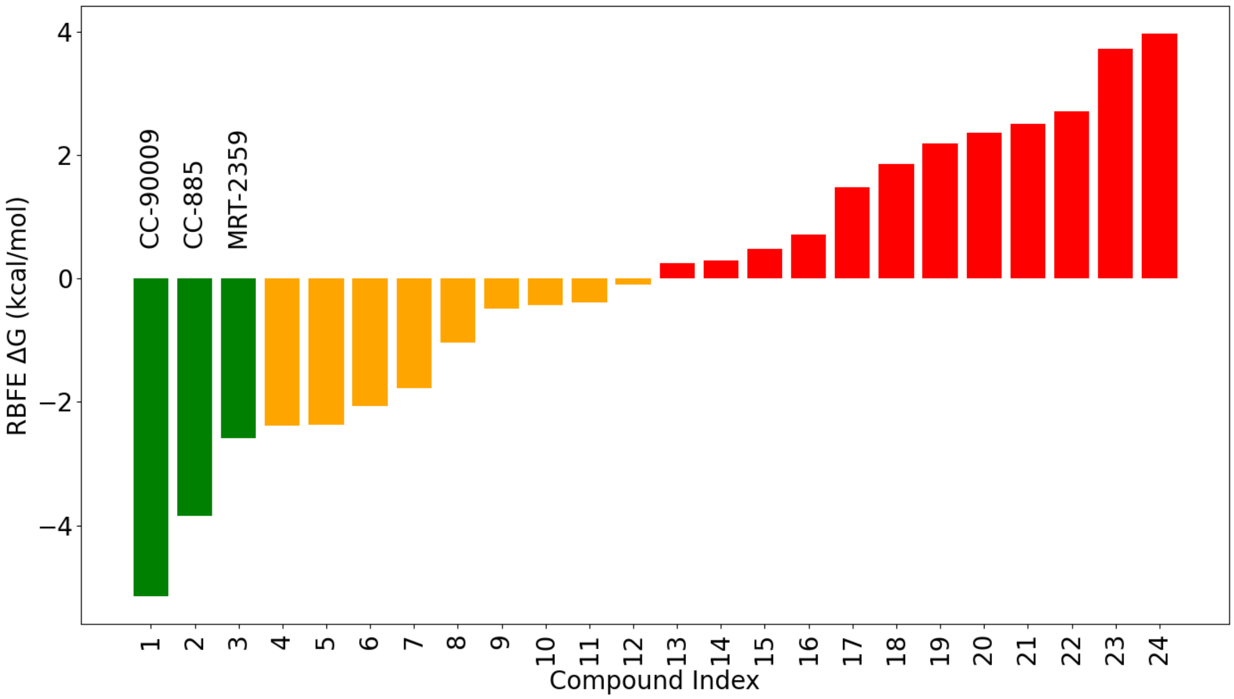


Figure S3: Molecular glues binding to GSPT1-CRBN complex. The three lowest values, corresponding to CC-90009, CC-885, and MRT-2359, are highlighted in green. Values less than zero are shown in orange, while values greater than zero are in red.

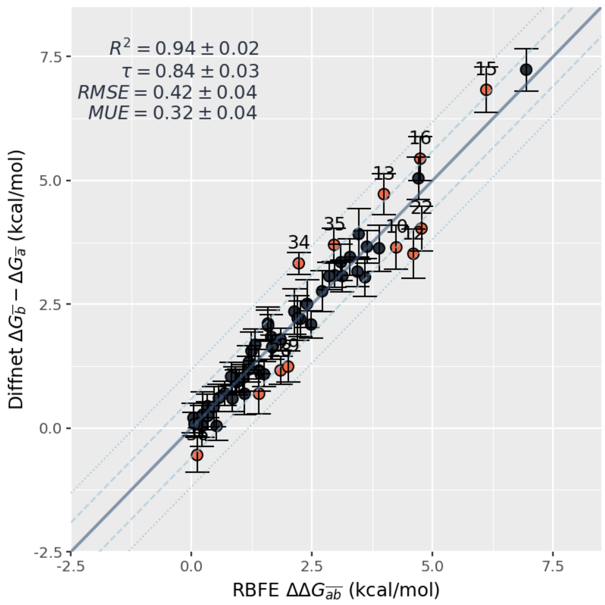


Figure S4: RBFE consistency plot of the 30 CRBN-GSPT1 compounds tested experimentally. The consistency plot illustrates the correlation between the $\Delta\Delta G_{RBFE}$ values, obtained from RBFE calculations, and from the DiffNet analysis of the $\Delta\Delta G_{RBFE}$ values. The dotted and dashed lines demarcate regions where the two predictions differ by $k_BT$ and $2k_BT$, respectively. The direction of each RBFE calculation is adjusted such that all plotted values are positive in the $x$-axis (indicated by the $\bar{a}, \bar{b}, \bar{ab}$ subscripts). $R^2$, $\tau$, $RMSE$, and $MUE$ denote, respectively, the correlation coefficient, Kendall rank correlation coefficient, Root Mean Square Error, and Mean Unsigned Error.

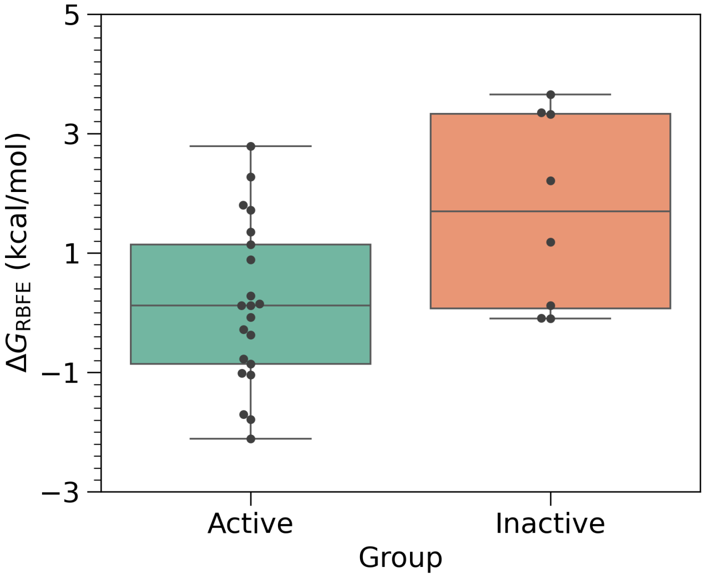


Figure S5: $\Delta G_{RBFE}$ distributions of the 30 CRBN-GSPT1 compounds tested experimentally. The difference between the average $\Delta G_{RBFE}$ of the group of active (n=22) and inactive (n=8) compounds is statistically significant as assessed by a two-tailed t-test ($p = 0.03$). The list of compounds is provided in Table S1.

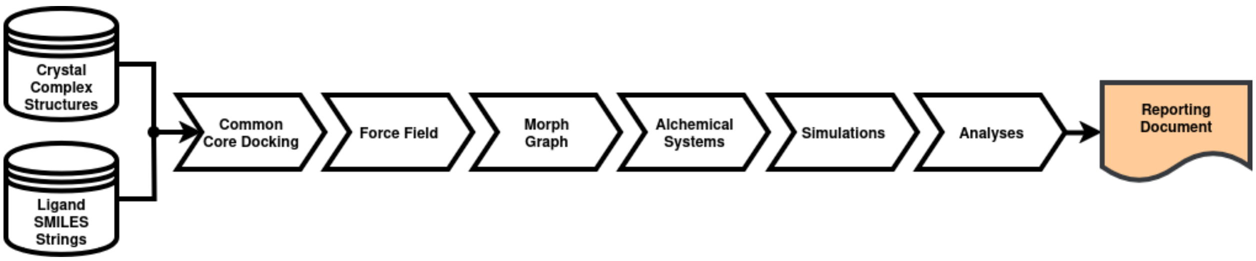


Figure S6: A flowchart showing the RBFE workflow.

Table S1: Experimentally tested molecular glues ordered by $DC_{50}$. The value $DC_{50} > 10000$ nM is assigned to inactive compounds, as 10000 nM was the highest compound concentration tested in our assay. The catalog id can be used to purchase compounds from either the Enamine REAL or WuXi Galaxi libraries.

| Name | $DC_{50}$ (nM) | | Catalog ID |
|---|---|---|---|
| | $n = 1$ | $n = 2$ | |
| MG15 | 7.2 | 11.3 | m_2708bbb____23836964____24002194 |
| MG6 | 8.0 | 5.9 | m_2708bbb____23836964____6201402 |
| MG8 | 12.2 | 14.2 | m_2708bbb____23836964____2680170 |
| MG25 | 13.0 | 15.0 | m_2708bbb____23836964____1324062 |
| MG3 | 15.7 | 17.6 | m_2430bbb____16899452____20031314 |
| MG1 | 16.6 | 22.0 | m_2430bbb____881166____20031314 |
| MG21 | 21.3 | 37.3 | m_2708bbb____23836964____17558532 |
| MG26 | 24.4 | 37.3 | m_2708bbb____23836964____915200 |
| MG29 | 26.8 | 26.2 | m_2708bbb____23836964____23985738 |
| MG27 | 34.7 | 89.4 | m_2708bbb____23836964____10126418 |
| MG13 | 41.4 | 70.8 | m_2708bbb____23836964____13778296 |
| MG9 | 48.3 | 74.9 | m_11bbb____20031326____14417494 |
| MG19 | 62.3 | 91.5 | m_2708bbb____23836964____5359764 |
| MG7 | 63.4 | 171.0 | m_2708bbb____23836964____24023478 |
| MG18 | 95.5 | 132.0 | m_22bbn____765804____26254764 |
| MG2 | 199.0 | 480.0 | m_2708bbb____23836964____6406720 |
| MG28 | 255.0 | 666.0 | m_2430bbb____12213720____20031314 |
| MG12 | 567.0 | 1070 | m_2708bbb____23836964____914558 |
| MG23 | 635.0 | 1230 | m_2708bbb____23836964____23999170 |
| MG20 | 690.0 | 2580 | m_2708bbb____23836964____23985336 |
| MG11 | 735.0 | 1540 | m_2708cbb____23836964____5532466 |
| MG17 | 1410 | 3190 | m_2708bbb____23836964____23985596 |
| MG16 | >10000 | >10000 | m_2708bbb____23836964____913200 |
| MG5 | >10000 | >10000 | m_2430bbb____3114690____20031314 |
| MG22 | >10000 | >10000 | m_2708bbb____23836964____23995280 |
| MG24 | >10000 | >10000 | m_2708bbb____23836964____24000588 |
| MG10 | >10000 | >10000 | m_11bbb____20031326____25928494 |
| MG4 | >10000 | >10000 | m_2708bbb____23836964____913562 |
| MG14 | >10000 | >10000 | m_2430bbb____22526182____20031314 |
| MG30 | >10000 | >10000 | m_2430bbb____22529126____20031314 |

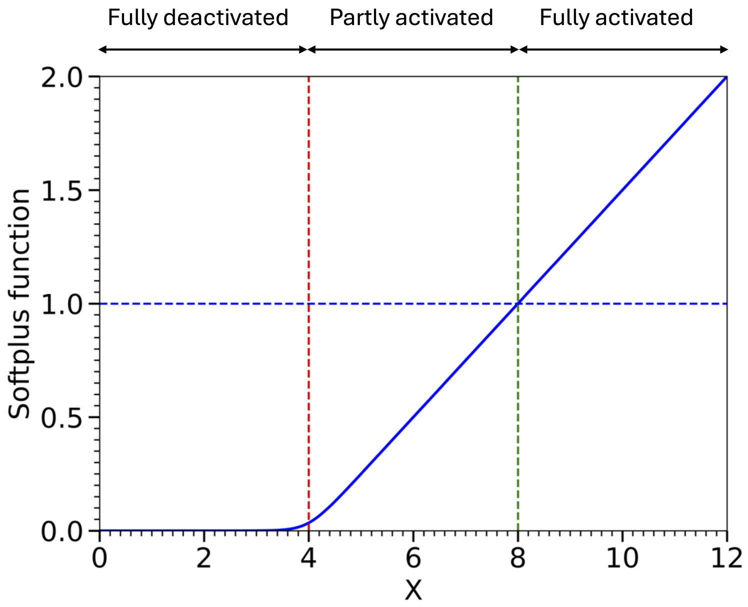


Figure S7: Softplus function $f(X) = 0.05 \cdot \ln\left(1 + \exp^{20 \cdot \frac{X - Xa}{Xb - Xa}}\right)$ (continuous blue line) with $Xa = 4$ and $Xb = 8$. The dashed red, green and blue lines represent $Xa$, $Xb$ and a cutoff value. Together they determine the three regions of activation of the softplus function.
